# Impact of early insect herbivory on the invasive oak lace bug (*Corythucha arcuata* Say, 1832) in different oak species

**DOI:** 10.1101/2022.11.30.518479

**Authors:** Elena Valdés-Correcher, Maarten de Groot, Laura Schillé, Alex Stemmelen, Yannick Mellerin, Olivier Bonnard, Bastien Castagneyrol

## Abstract

Insect herbivores co-occurring on the same host plant interact in various ways. In particular, early-season insect herbivory triggers a wide range of plant responses that can determine the performance of herbivores colonizing the plant later in the course of the season. But the strength and direction of such effects are debated, and virtually unknown in the case of novel interactions involving exotic insects in their introduction range. We conducted an observational field study in SW France, a region recently invaded by the Oak Lace Bug (OLB, *Corythucha arcuata* Say). We measured early chewing damage and subsequent OLB damage in four oak species (*Quercus robur, Q. pubescens, Q. cerris* and *Q. ilex*). We set up a complementary non-choice experiment in the laboratory, feeding OLB with leaves with or without prior herbivory. The four oak species differed in their sensitivity to OLB damage, *Q. ilex* being broadly resistant. Prior herbivory promoted OLB damage in the laboratory experiment, but not in the field. However, prior herbivory did not alter the rank of oak resistance to the OLB. Our results suggest possible synergistic effects between spring defoliators and the OLB. This study brings insight into herbivore-herbivore interactions and their possible implications for forest management.

## Introduction

Plants have long been seen as providing insect herbivores with such a large abundance of resources and niches that interactions among herbivores co-occurring on the same plant were deemed unlikely (reviewed by Kaplan and Denno, 2007). Yet, advances in our understanding of the mechanisms of plant direct and indirect defenses against herbivores largely challenged this view. It is now well established that insect herbivores co-occurring on the same host plants, either simultaneously, during the course of a growing season or even beyond, do interact among each others in various ways (Poelman et al. 2008; Moreira et al. 2018; Fernández de Bobadilla et al. 2021), regardless of whether they belong to the same species or not (Kaplan and Denno 2007; Van Dijk et al. 2020; Castagneyrol et al. 2021). In particular, early-season insect herbivory triggers a wide range of plant responses, ranging from induced defenses (Thaler et al. 2012; Moreira et al. 2018; Abdala-Roberts et al. 2019) to altered nutritional quality (Newingham et al. 2007; Gómez et al. 2010, 2012; Moreira et al. 2012). These changes indirectly influence and determine the performance of subsequent herbivores (McArt and Thaler 2013; Moreira et al. 2015), as well as the structure of community of species associated with the plant (Poelman and Dicke 2014; Biere and Goverse 2016). However, how early-season herbivory influences late season herbivory is paradoxically not well known.

The effect of early insect herbivory on subsequent herbivory varies substantially among insect herbivores and feeding guilds (Bingham and Agrawal 2010; Carmona and Fornoni 2013; Moreira et al. 2013; Schaeffer et al. 2018) and ranges from negative to positive (Kaplan and Denno 2007; Ohgushi 2008). Plant response to biotic aggressions is coordinated by two antagonistic signaling pathways (Erb et al. 2012). Plant pathogens or sucking insects triggers the Salicylic Acid (SA) defense pathway (Leitner et al. 2005; Schweiger et al. 2014), whereas necrotrophic pathogens, chewing or mining herbivores trigger the Jasmonic Acid (JA) defense pathway (Glazebrook 2005; Thaler et al. 2012). The reciprocal antagonistic interactions between the SA and JA pathways are responsible for negative interactions between herbivores belonging to the same guild, and positive interactions between herbivores of different guilds when sequentially attacking the same plant individual (Moreira et al. 2018). For instance, Rigsby et al., (2021) study showed that the positive effect of previous hemlock wolly adelgid (*Adelge tsugae*) on later-instar spongy moth (*Lymantria dispar*) attack on eastern hemlock (*Tsuga canadiensis*) was mediated by an increase on SA pathway and consequent reduction in JA pathway as a result of hemlock wolly adelgid attack. However, most of previous studies have been performed under experimental conditions, making it difficult to assess the relative importance of this phenomenon in the real world (but see Hernandez-Cumplido et al., 2016; Kinahan et al., 2020).

Biological invasions create new biotic interactions between insects and their host plants, as well as among insects co-occurring on the same plant (Kinahan et al. 2020; Rigsby et al. 2021). The introduction of insect herbivores in new areas is a major cause of disturbance in natural and managed ecosystems. For instance, the invasive pest emerald ash borer (*Agrilus planipennis*) has killed millions of ash trees (*Fraxinus* spp.) in North America (Herms and McCullough 2014) causing cascading direct and indirect effects on forest community composition and ecosystem processes (Gandhi and Herms 2010). If indirect interactions between herbivores exploiting the same resource are a widespread phenomenon, the damage caused by exotic herbivores may accumulate or, on the contrary, reduce the damage caused by native herbivorous insects. The invasive oak lace bug (OLB, *Corythucha arcuata* Say), a native species from North America, is a recent example of a fast-spreading non-native insect with considerable damage capability in its recently invaded area of distribution (Paulin et al. 2020). By feeding on oak leaves, OLB also interacts with other insect herbivores (e.g. chewers and leaf galls; Paulin et al., 2019) and native natural enemies (e.g. coccinellids, predatory bugs and spiders; Paulin et al., 2020). Previous studies have investigated the effect of OLB on insect herbivores and have shown that OLB has a negative effect on the development of some leaf galling species and chewer herbivore species (Paulin et al. 2019). The other way around, whereby the OLB is influenced by the resident herbivore community, is however unknown. Being a multivoltine sucking insects that triggers the SA defense pathway (Leitner et al. 2005; Schweiger et al. 2014), theory predicts that the development of the OLB in the course of the season is facilitated by early attacks of its hosts by insects triggering the JA defense pathway, typically defoliators (reviewed by Moreira et al., 2018). Investigating the effect of early insect herbivory on late OLB attacks may help us to better understand the mechanisms that allow this invasive species to build and maintain outbreak densities.

In this study, we investigated the effect of prior chewing herbivory on oak lace bug (OLB) damage and mortality, combining field and laboratory experiments. To that aim, we measured chewing damage early in the season and OLB damage late in the season on fifteen trees of four oak species (*Q. robur, Q. pubescens, Q. cerris* and *Q. ilex*) in a newly invaded area. We complemented this observational field study with a controlled no-choice experiment in the laboratory, assessing subsequent OLB damage and mortality on intact *vs* damaged leaves of the same four oak species. We predicted that (1) early-season chewing damage has a positive effect on subsequent OLB damage, OLB abundance and OLB eggs clutches abundance, and that (2) these effects vary among oak species in natural conditions. We also predicted that (3) early chewing damage favor posterior OLB damage and reduces its mortality, and that these effects vary among oak species under controlled laboratory conditions.

## Material and methods

### Target species

The OLB is a herbivore native to North America. It was introduced in Europe and first found in Italy and Turkey in 2000. It is thought that the Balkan and Central European population was spread from Turkey (Bernardinelli 2000; Csóka et al. 2020; Paulin et al. 2020) and that likely it was spread from Italy to France. The OLB mainly feeds on species of the *Quercus* genus, most frequently attacking *Q. robur* (L.), *Q. petraea* (Matt.) Liebl, *Q. frainetto* (Ten.) and *Q. cerris* (L.) (Csóka et al. 2020) but also other species such as *Castanea sativa* (Mill.) and *Rosa canina* (Bernardinelli 2000). This wide variability of suitable hosts allows its expansion in Europe and Asia (Csóka et al. 2020; Paulin et al. 2021). It reaches high densities late in the season (late July, late August and September), frequently covering the integrity of oak leaves and causing chlorosis, discoloration and desiccation of the leaf surface, reducing the photosynthesis and even causing premature leaf fall and acorn abscission (Dobreva et al. 2013; Paulin et al. 2020). The OLB was first observed in the study area in 2018 (B. Castagneyrol, personal observation).

### Observatory field study

#### Study site

The field experiment was carried out in the city of Bordeaux, SW France, a region characterized by an oceanic climate with mean annual temperature of 12.8 °C and annual precipitation of 873 mm over the last 20 years. We selected four oak species – namely the pedunculate (*Quercus robur*), pubescent (*Q. pubescens*), Turkey (*Q. cerris*) and holm (*Q. ilex*) oaks – in five parks where the four species co-occurred (**Figure 1**).

**Figure 1.**
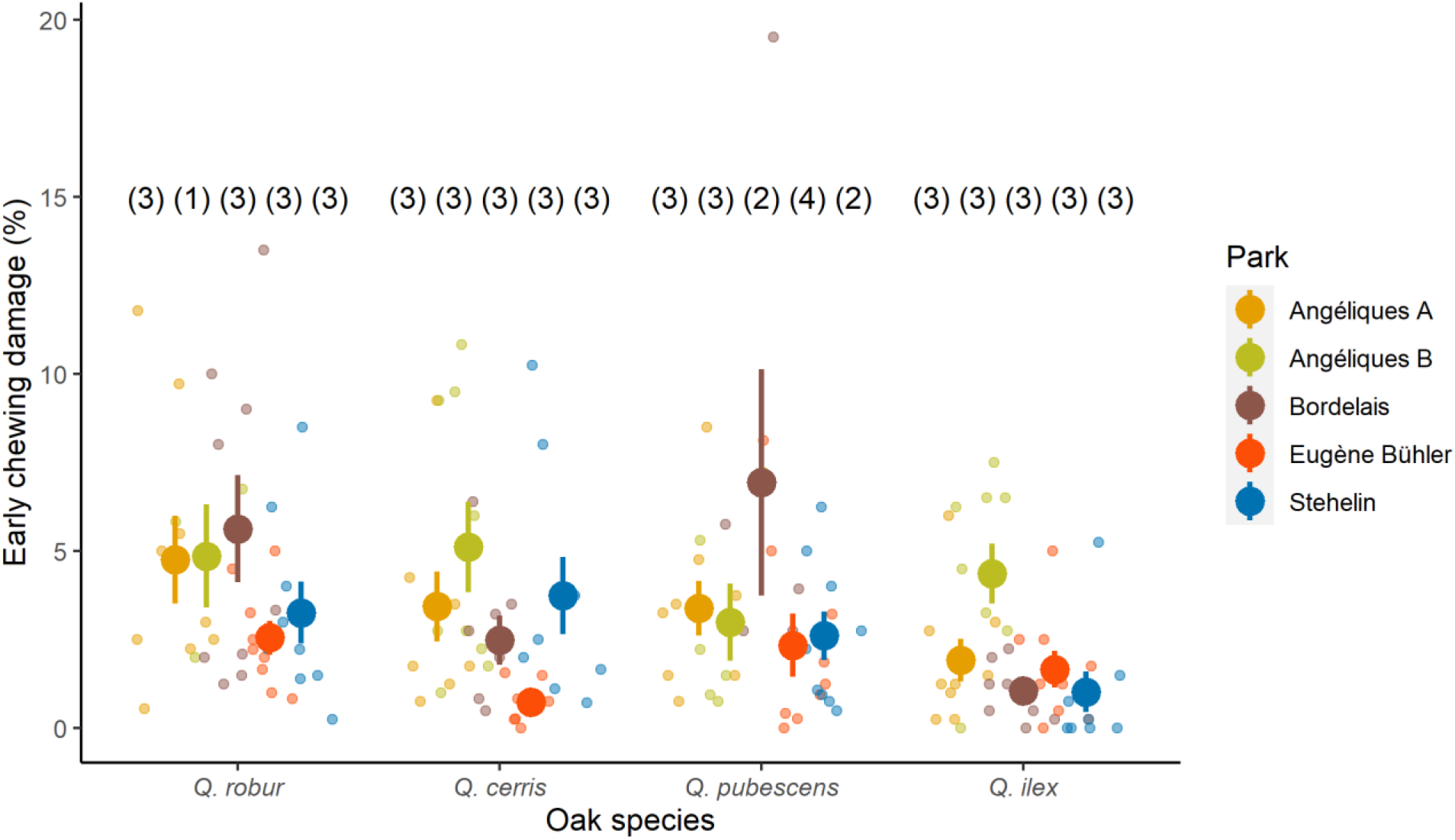
Percentage of early chewing damage averaged at branch level in each park and oak species. Numbers in brackets correspond to the number of trees selected in each park and oak species. Large solid dots and error bars represent raw means ± SE; small dots represent raw data.

We haphazardly selected one to four mature oak trees per species provided with low hanging branches that could be reached from the ground (57 oaks) in Angéliques A (44°50N 0°33’W), Angéliques B (44°51N 0°33’W), Eugène Bühler (44°52’N 0°34’W), Stehelin (44°51’N 0°37’W) and Bordelais (44°51’N 0°36’W) parks (**Figure 1**). The selected species represent 70 % of the oak species present in the public domain in Bordeaux city. All but *Q. cerris* are native to the region. A preliminary survey conducted in 2021 on the 10 most abundant oak species present in the public domain of Bordeaux city found that more than 80% of *Q. robur* (n = 169 trees surveyed), 85% of *Q. pubescens* (n = 269) and 60% of *Q. cerris* (n = 402) were attacked by the OLB, whereas less than 5% of *Q. ilex* (n = 168) were attacked (B. Castagneyrol and A. Stemmelen, unpublished data). Host range in the study area was therefore consistent with that observed in other countries where the OLB was introduced a decade ago (Csóka et al., 2020).

#### Leaf herbivory

In May 2022, we haphazardly selected three branches facing opposite directions in each tree. In each branch, we haphazardly selected one leaf, and attached a thin metallic wire at its basis. We then tagged every leaf along the branch, going down to the branch basis, up to 10 leaves. The first and fifth leaves had metallic wires of different colors from leaves 2-4 and 6-10. In total, we identified 30 leaves per tree and 450 leaves per species.

We took a high resolution pictures of each leaf in late May and late July to later quantify early chewing damage (% leaf area removed or impacted by chewing and mining herbivores) and subsequent OLB damage (% leaf area with chlorosis) as well as the number of OLB individuals and egg clutches. Early chewing damage and subsequent OLB damage were visually scored by assigning each leaf to one of the following classes: 0, 0.1–5.0, 5.1–10.0, 10.1–15.0, 15.1–25.0, 25.1–50.0, 50.1–75.0 or >75%. We then used the midpoint of each class to average both chewing and OLB damage at the branch level, and calculated the mean number of OLB insects and egg clutches per branch. Early OLB damage was observed in no more than 11% of leaves in the first survey and was not considered in further analyses.

We found a positive correlation between OLB damage and the abundance of OLB individuals (Spearman’s r = 0.83) and egg clutches abundance (r = 0.82). The results of the analyses were qualitatively the same with the three response variables. We decided to only present data on OLB damage in the main text, for it is the most comparable between the field and laboratory study. The results of the analyses on the number of OLB individuals and egg clutches are available in the supplementary material (**Table S1**, **Figure S1**).

### Laboratory experiment

We conducted a no-choice experiment in the laboratory in June 2022 with *Q. robur*, *Q. pubescens*, *Q. cerris* and *Q. ilex* leaves sampled in one of the four parks used in the field study (Park Angéliques A). We haphazardly sampled from three to five 50 cm branches on five trees per species and immediately brought them to the laboratory.

We haphazardly collected 10 damaged and 10 intact leaves per tree, and then pooled leaves from the different branches, keeping track of tree identity. We thus constituted a random pool of 50 damaged and 50 intact leaves per species, i.e., 400 leaves. We ensured no chlorotic spots or OLB individuals were present on sampled leaves.

We prepared a total of 400 petri dishes, each receiving one intact or damaged leaf from one of the four oak species and one OLB nymph. We collected nymphs on the same branches, the same day we sampled branches. We placed a piece of wet filter paper at the bottom to provide the necessary moisture to maintain the leaves and nymphs in optimal conditions. We stored petri dishes in 50 hermetic plastic boxes filled with a thin layer of water to maintain relative humidity. Each box contained eight petri dishes consisting of a replica of intact and damaged leaves of each oak species. The experiment was conducted at room temperature (~20 °C) and lasted for eight days after which we recorded OLB mortality and damage (% leaf area with chlorotic spots). We used the same percentage classes as described above and averaged OLB damage at tree level within each level of early chewing damage. We excluded six leaves (one from *Q. cerris*, one from *Q. pubescens* and four from *Q. robur*) on which it was not possible to measure OLB damage because they were covered with fungus. Thus, a total of 394 leaves were surveyed.

### Statistical analyses

Preliminary data exploration confirmed that *Q. ilex* was a poor host species for the OLB: none of the 450 leaves sampled in the field had chlorotic spots, which were only found in 2 leaves (2%) in the no-choice experiment. We first ran non-parametric Kruskal-Wallis tests followed by pairwise comparisons to test differences in OLB damage among the four species. We restricted further analyses to data collected in the three other species (*Q. robur*, *Q. pubescens* and *Q. cerris*).

#### Field survey

First, we tested the effects of early chewing damage and oak species as well as their interaction on OLB damage, OLB insect and egg clutches abundance at branch level, in natural conditions with linear mixed effect models (LMM). Early season chewing damage (continuous variable) and oak species (three-levels factor) were included as fixed effects, and Tree ID as random effect nested within Park ID to account for pseudo-replications of branches within trees and trees within parks at each park and tree. Some leaves were not retrieved during the second survey and were therefore not considered in the analyses. The final dataset represented a total of 42 trees with 1106 leaves and 119 branches.

#### Laboratory no-choice experiment

We tested the effect of early chewing damage (two-levels factor: intact *vs* damaged) and oak species (three-levels factor) on OLB damage and mortality in separate generalized mixed-effects models (GLMM). The OLB damage model had a Gausian error distribution whereas the mortality model has a binomial error distribution and logit-link. Early chewing damage and oak species were included as fixed effects and Tree ID as random effect to account for repeated measurements on each tree. We averaged OLB damage and summarized dead OLB *vs* alive OLB per tree (total: *n* = 15 trees, excluding *Q. ilex* leaves).

We estimated and compared (G)LMM fit by calculating marginal and conditional *R^2^* (respectively *R^2^m* and *R^2^c*) in order to estimate the proportion of variance explained by fixed (*R^2^m*) and fixed plus random factors (*R^2^c*) (Nakagawa and Schielzeth 2013). All analyses were conducted in the R 4.0.5 (R Core Team 2020). We ran (G)LMM and calculated summary statistics and model coefficients with functions provided in libraries with package *lme4* (*lmer* and *glmer* functions) (Bates et al. 2018).

## Results

### Field experiment

Early chewing damage represented on average (± SE) 3.6 ± 0.21% of the total leaf area early in the season, while OLB damage represented on average 11.4 ± 0.51% of the leaf area (42 trees and 1106 leaves, excluding *Q. ilex* trees) (**Figure 1**). OLB abundance was on average 1.7 ± 0.21% of the inspected leaves, whereas OLB egg clutches abundance was on average 0.5 ± 0.03% of the inspected leaves late in the season.

OLB damage varied significantly among species (**Table 1**), with no OLB damage found in *Q. ilex*. Specifically, it was greater in *Q. robur* and *Q. cerris* than in *Q. pubescens* (**Figure 2**). OLB damage was also significantly lower on *Q. ilex* than on the rest of the oak species (Kruskal-wallis followed by post-hoc test). OLB damage was not influenced by early herbivory or by the interaction between oak species identity and early herbivory (**Table 1**).

**Figure 2.**
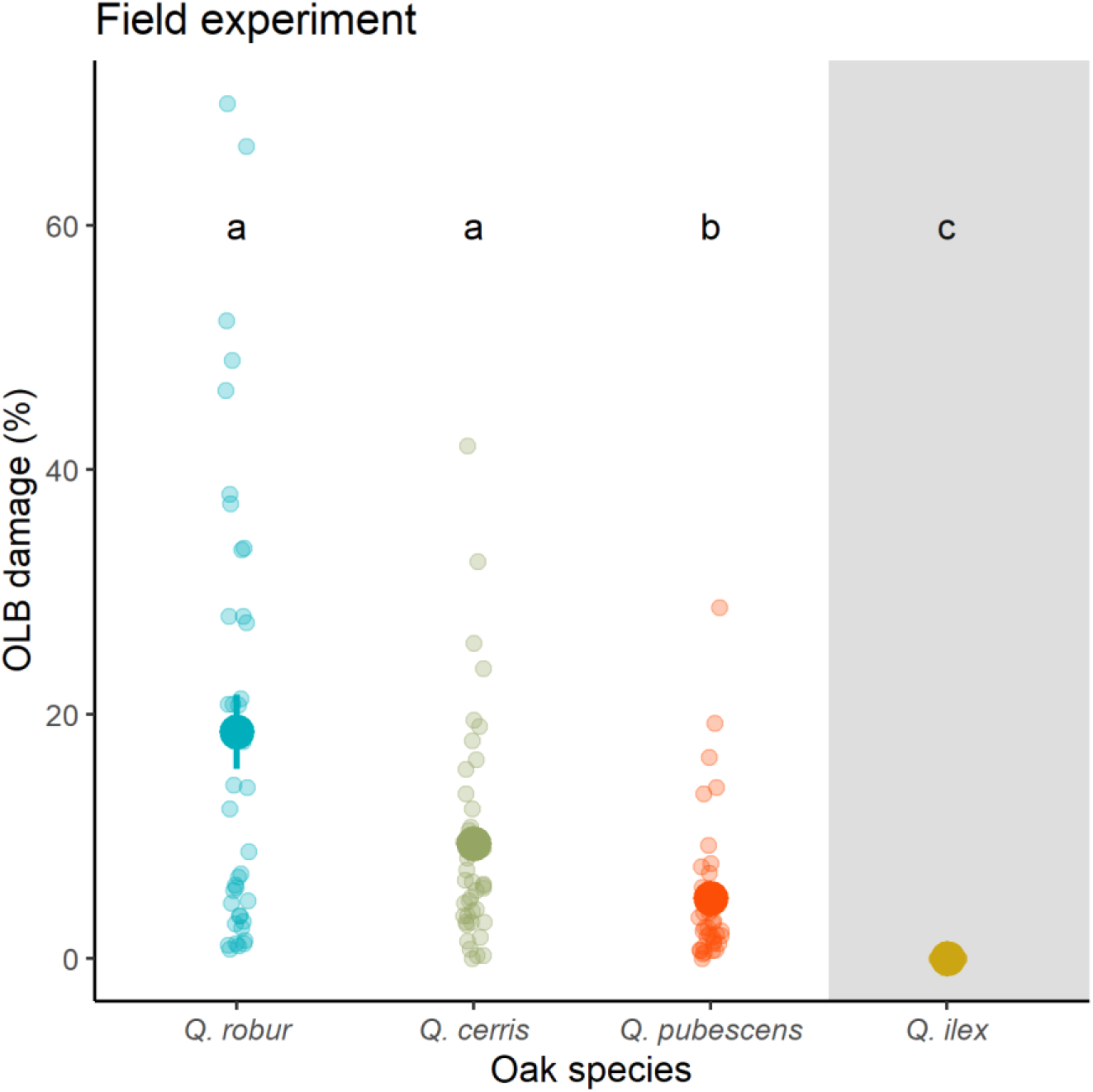
Variation of OLB damage (% chlorosis) among oak species in the field experiment. Small dots represent raw data, large symbols and error bars represent raw means ± SE. The gray shading area indicates that *Q. ilex* was not included in the LMM. Different letters (a, b and c) indicate significant differences between oak species obtained with the post-hoc test, and were the same as in the LMM with only the three other oak species

**Table 1.**
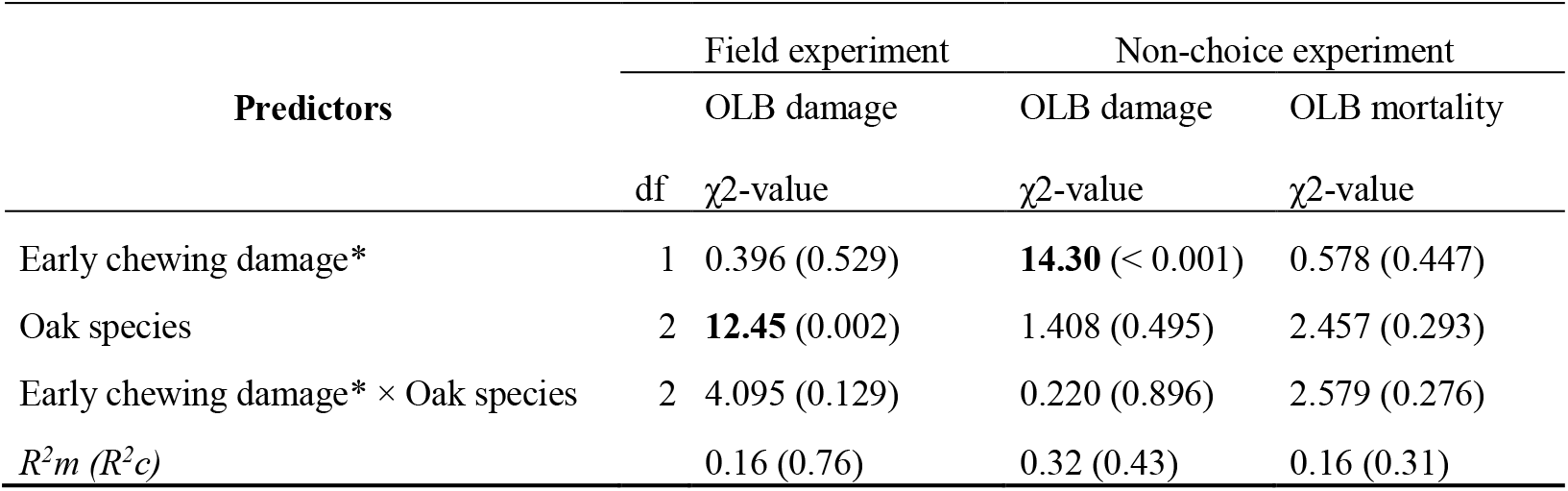
Summary of (G)LMMs testing the effect of early chewing damage*, oak species and their interaction on OLB damage and mortality. P-values are indicated within brackets and significant effects are in bold. Marginal (*R^2^m*) and conditional (*R^2^c*) *R^2^* are reported. *Early chewing damage is treated as a continuous and categorical variable in the field and laboratory experiments, respectively.

### Non-choice experiment

After eight days of the no-choice experiment, chlorotic spots caused by OLB covered 20.1 ± 100% of leaf surface. This raw mean was strongly influenced by the inclusion of *Q. ilex* leaves in the calculation and both OLB damage and mortality was significantly different between *Q. ilex* and the rest of oak species (Kruskal Wallis). Excluding this species, OLB damage was on average 26.5 ± 1.07%. Of the 392 OLB nymphs used in the experiment, 136 were dead after eight days (i.e., 34.7%), including 92 that were forced to feed on *Q. ilex* leaves (i.e., 23.5% of the total).

Contrary to what we observed in the field, OLB damage did not differ significantly among the three deciduous oak species (**Table 1, Figure 3**) but was significantly lower on *Q. ilex* than on the rest of the oak species (Kruskal-wallis followed by post-hoc test, **Figure S2**). It was on average 23.5% higher on leaves that had been previously attacked by chewing herbivores (**Table 1, Figure 3**). The effect of early herbivory on OLB damage was consistent across oak species (no significant *Herbivory* × *Species* interaction, **Table 1**). OLB mortality did not vary between herbivory treatments or among the three oak species (**Table 1**), but was significantly higher on *Q. ilex* than on the rest of the oak species ((Kruskal-wallis followed by post-hoc test, **Figure S2**).

**Figure 3.**
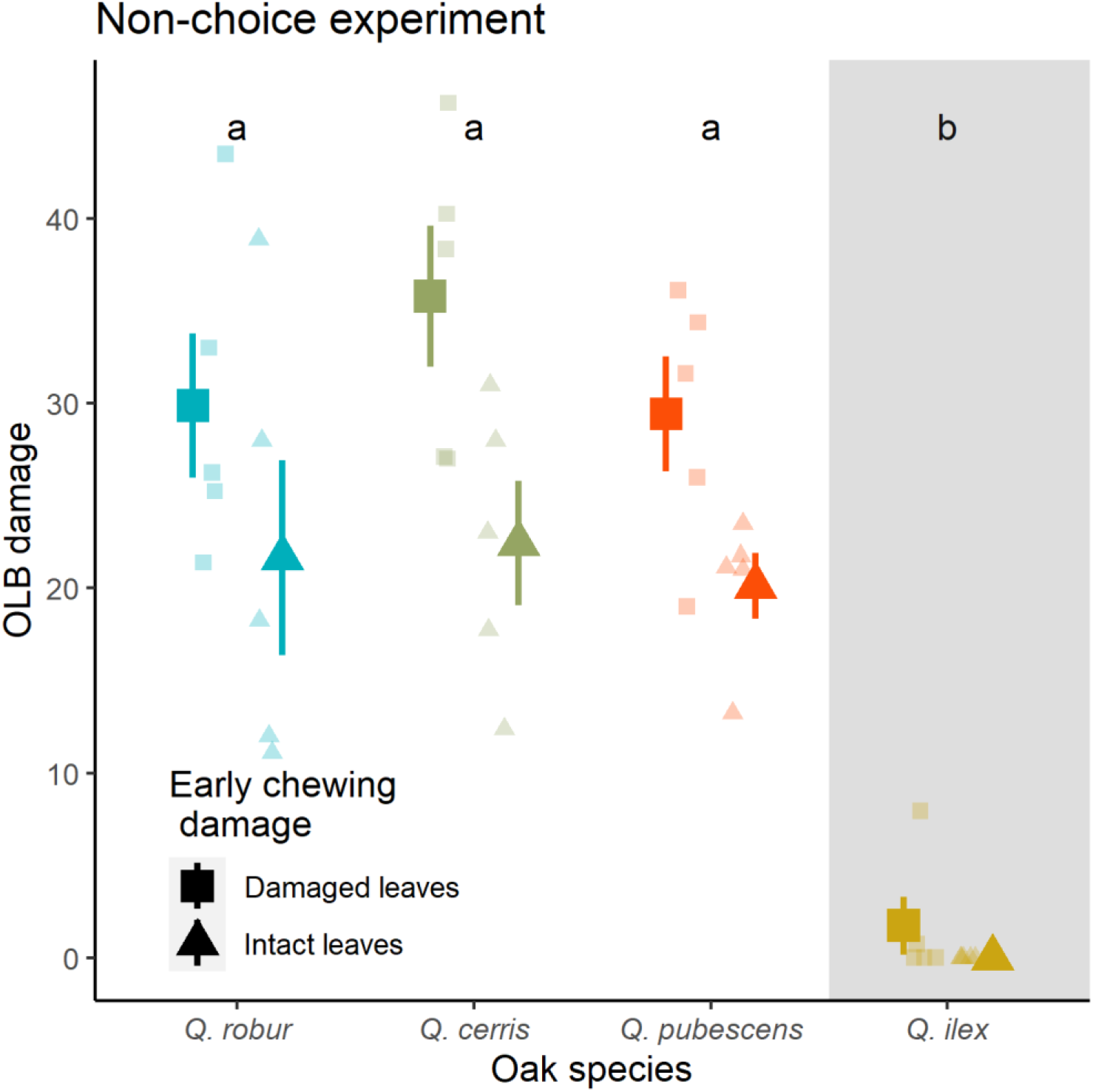
Variation of OLB damage (% chlorosis) among oak species in the non-choice experiment. Small symbols represent raw data, large symbols and error bars represent raw means ± SE. The gray shading area indicates that *Q. ilex* was not included in the GLMM. Different letters (a and b) indicate significant differences between oak species). Significant differences were obtained with the post-hoc test, and were the same as in the LMM with only the three other oak species

## Discussion

We found that subsequent damage caused by the invasive oak lace bug (OLB) *Corythucha arcuata* to three of its hosts in the European introduced range were facilitated by early chewing damage. This effect was however only observed under controlled laboratory conditions. On the contrary, we observed significant differences in host sensitivity to the OLB only in the field. We found no evidence that early herbivory influenced the rank of sensitivity of different host species to the OLB.

Early chewing damage increased leaf sensitivity to the OLB in a non-choice experiment but it did not influence OLB mortality as larvae were still able to feed on intact leaves. As the OLB is a piercing-sucking herbivore, this result is consistent with previous studies having reported an antagonism between the JA and SA pathways, the former being triggered by and directed to chewing herbivores, and the latest by and to sap-feeding herbivores (Thaler et al. 2012; Moreira et al. 2018). However, how much this cross-talk influences herbivore-herbivore interactions in the real world is controversial. For instance, positive interactions were documented between the hemlock woolly adelgid (*Adelge tsugae*) and the spongy moth (*Lymantria dispar*) (Kinahan et al. 2020). On the contrary, Ali and Agrawal (2014) found that although monarch larvae (*Danaus plexippus*) triggered the JA pathway in damaged milkweed (*Asclepias syriaca*), the subsequent downregulation of the SA pathway had no noticeable effects on the oleander aphid (*Aphis nerii*). The fact that the facilitative effect of prior herbivory on OLB damage was only observed under controlled laboratory conditions further questions the mechanisms at play and their relative importance.

It is true that a non-choice experiment may not be representative of what occurs in the wild where herbivores move freely and can choose their food based on its perceived quality. But should have the OLB been able to select among damaged *vs* undamaged leaves, undamaged leaves would have been avoided and damaged leaves even more damaged than what we observed. In this case, it is particularly surprising that this pattern was not observed in the field. This suggests that other ecological factors with prominent importance determined the distribution of OLB and associated damage within oak foliage. Herbivores chose their feeding resource based on perceived quality (Mattson 1980; Wetzel et al. 2016), but also based on perceived risk (Jeffries and Lawton 1984). Thus, insect herbivores may choose plants that provide enemy-free space over the best food (Singer et al. 2004; Ghosh et al. 2022). It is also possible that the microclimate played a disproportionate role on the feeding behaviour of the OLB in the field, as has already been described for other species (Bernaschini et al. 2020; Rytteri et al. 2021; Stewart et al. 2021). This does not imply that prior herbivory was not influential, but merely that its effect was overcome by something else. In particular, the gregarious behaviour of the OLB lay may also alter the effect of previous herbivory. For instance, Kroes et al. study (2015) showed that cabbage aphids (*Brevicoryne brassicae*) manipulate defense induction in the thale cress (*Arabidopsis thaliana*), in a density-dependent manner. It is also possible that predation altered herbivore-herbivore interactions in the field, which remains difficult to assess given the very little knowledge we have on OLB enemies in its introduced range (Bernardinelli and Zandigiacomo 2000; Sönmez et al. 2016; Kovač et al. 2020; Paulin et al. 2020).

We revealed differences in oak sensitivity to the OLB that were independent of prior chewing damage. We found that OLB damage varied among oak species in the field experiment, but not in the non-choice experiment. These results may indicate that OLB damage rate can be similar on *Q. robur*, *Q. cerris* and *Q. pubescens* in situations when only confronted by one of these host species as was also shown in the non-choice experiment. In situations when OLB can choose, like in the field study, OLB shows preferences for specific oak species such as *Q. robur* and *Q. cerris*. More specifically, OLB damage was significantly higher on both *Q. robur* and *Q. cerris* than on *Q. pubescens* in the field experiment, suggesting that both *Q. robur* and *Q. cerris* are more susceptible and are more suitable hosts to OLB damage. Similar differences in insect herbivore damage among oak species have been also described on OLB (Marković et al. 2021) as well as on other insect herbivore species such as *Lymantria dispar* and *Thaumetopoea processionea* (Milanović et al. 2014; Damestoy et al. 2019, 2021). It is however important to recall that preferences for a given species as measured by host choice by mobile herbivore stages may not perfectly reflect the subsequent amount of damage (Marković et al. 2021). Still, it is noticeable that, overall, the hierarchy of oak sensitivity to the OLB was consistent across studies, thus calling for further examination of defense traits at play.

Despite positive interactions between defoliators and OLB, prior herbivory did not make *Q. ilex* a more suitable host. Similarly, Csóka et al., (2020) study also found that *Q. ilex* leaves are not attacked by OLB in any of the forest stands recorded in ten countries (Bosnia and Herzegovina, Bulgaria, Croatia, Hungary, Italy, Romania, Russia, Serbia, Slovenia and Turkey). This deciduous oak species is mainly attacked by fungal pathogens such as *Phytophthora quercina*, *P. cinnamomi* and *P. ramorum*; as well as by polyphagous lepidopterans such as *Lymantria monacha*, *L. dispar*, *Tortrix viridana* and *Malacosoma neustria* (De Rigo and Caudullo 2016). However, some of the characteristics of its leaves, such as their toughness, may not be attracted to sucking insects such as OLB, or even increases its mortality as it is the case of *Q. suber* (Bernardinelli 2006) and *Q. ilex* (Bernardinelli, 2006; Csóka et al., 2020, our results).

## Conclusion and implications

The facilitation of OLB damage by early chewing damage found in this study could indicate us that there might be a variability in the risk of OLB damage among populations depending on the diversity and activity of insect herbivores. As a consequence, synergistic effects between major spring defoliators such as the spongy moth (*Lymantria dispar*) and the OLB can take place and result in profound consequences on the structure of ecological communities-both trees and insects - and the dynamic of forest ecosystems in the study area. It is therefore of great importance to continue studying the interaction of this invasive species with other organisms and with the environment, as well as to study the consequences it may have to the plant growth and fitness of the different oak species that it attacks before its damage begins to be irreversible. These findings indicate that a large proportion of European deciduous forests may be particularly prone to damage by this invasive pest, which should be a greater concern for foresters that it is yet (Bălăcenoiu et al. 2021).

## Acknowledgments

The authors warmly thank Cloée Jean and Timothé Lajubertie for their technical assistance in the field and in the laboratory. M.G. did this study through the research core group “Forest biology, ecology and technology” (P4-0107) financed via the Slovenian Research Agency.

## Rights and permissions

For the purpose of Open Access, a CC-BY 4.0 public copyright licence (https://creativecommons.org/licenses/by/4.0/) has been applied by the authors to the present document and will be applied to all subsequent versions up to the Author Accepted Manuscript arising from this submission.

## Author contributions

E.V.C, B.C, M.G, L.S and A.S contributed to the study conception and design. Material preparation and data collection were performed by E.V.C, L.S, O.B and Y.M. EVC estimated early chewing damage and OLB damage, while E.V.C, O.B and Y.M measured the OLB abundance and OLB egg clutches abundance. Data analysis was performed by E.V.C and BC. The first draft of the manuscript was written by E.V.C and all authors commented on previous versions of the manuscript. All authors read and approved the final manuscript.

## Data availability statement

The datasets generated during the current study are available from the corresponding author on reasonable request

## Supplementary Information

**Table S1.**
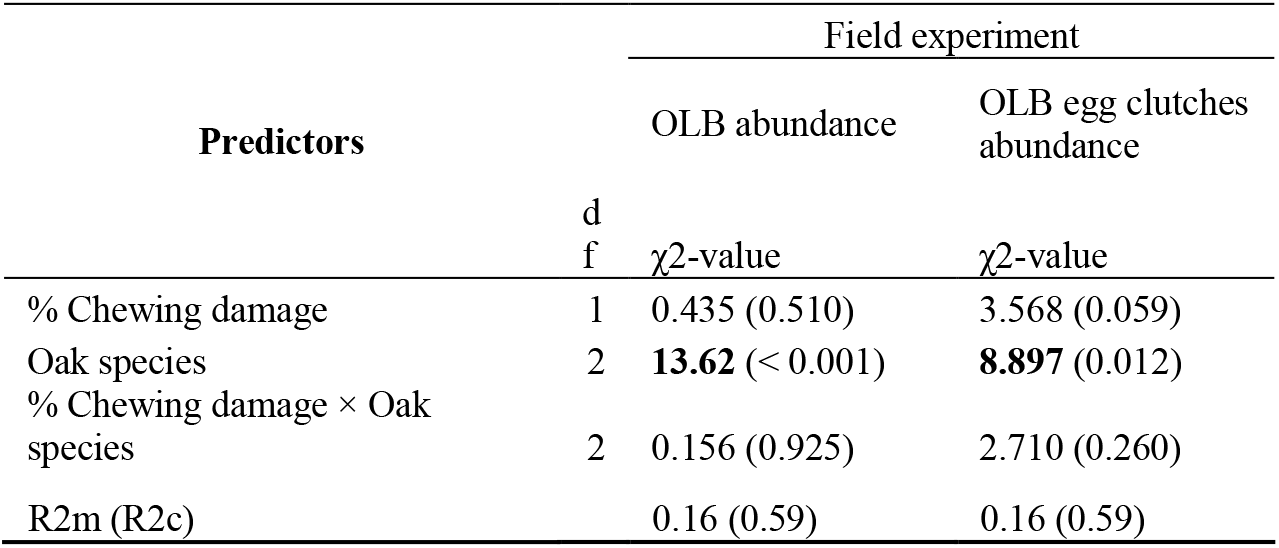
Summary of LMMs testing the effect of early chewing damage (% of chewing damage for the field experiment and oak species on OLB abundance and OLB egg clutches abundance. P-values are indicated within brackets and significant effects are in bold. Marginal (R^2^m) and conditional (R^2^c) R^2^ are reported.

**Figure S1.**
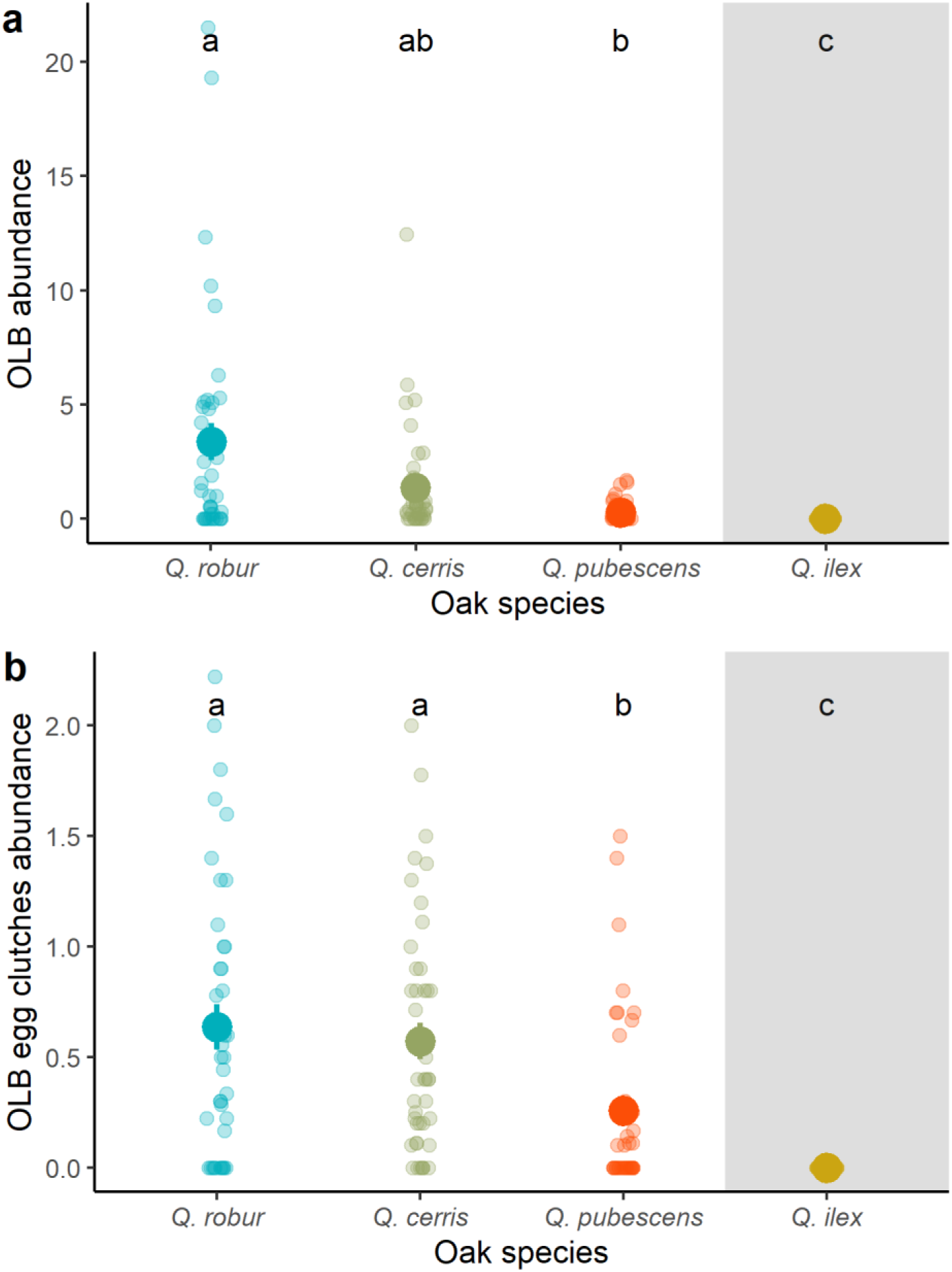
Variation of early OLB abundance (a) and OLB egg clutches abundance (b) among oak species in the field experiment. Small dots represent raw data, large symbols and error bars represent raw means ± SE. The gray shading area indicates that *Q. ilex* was not included in the LMM. Different letters (a, b and c) indicate significant differences between oak species obtained with the post-hoc test and were the same as in the LMM with only the three other oak species

**Figure S2.**
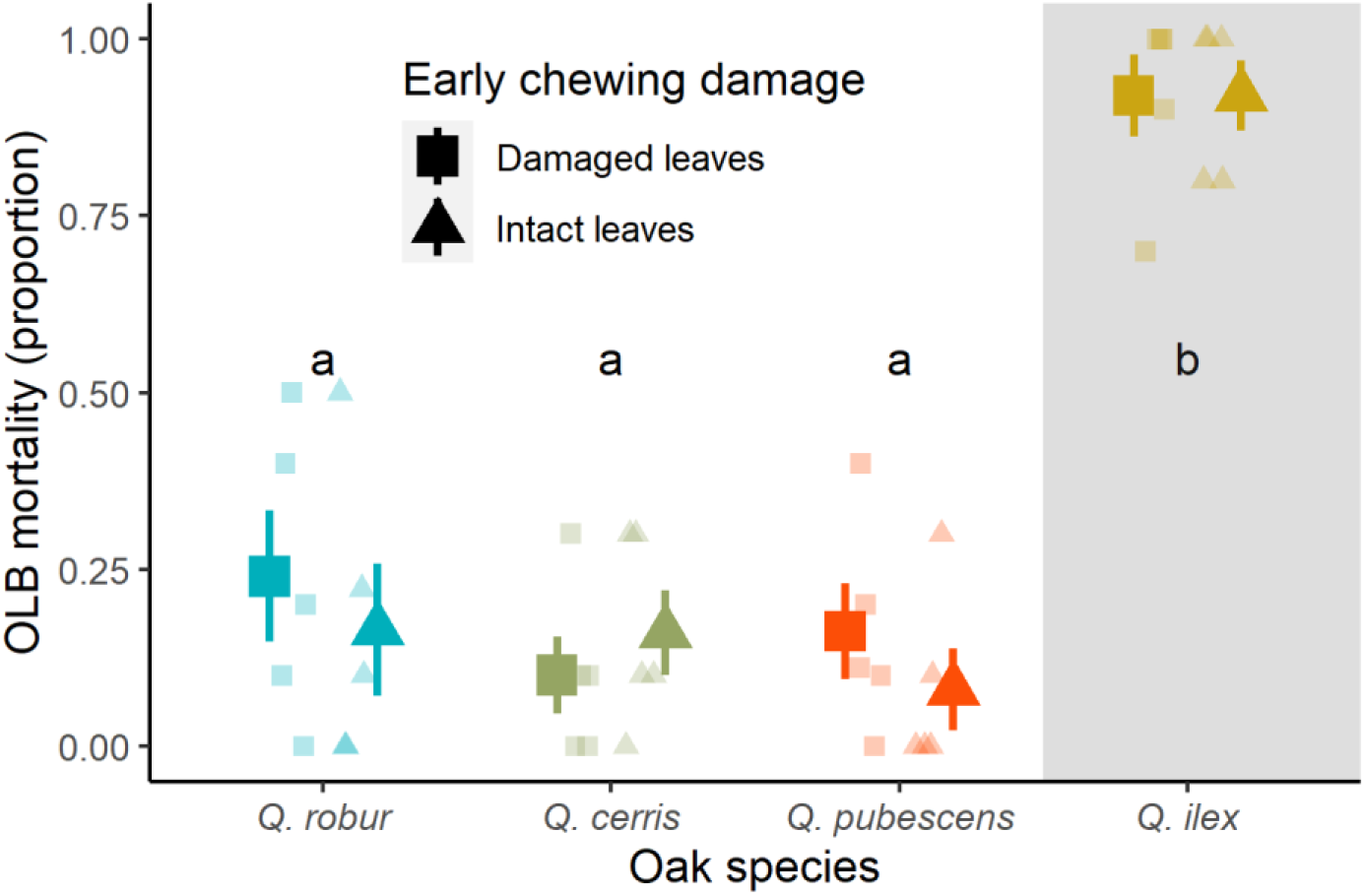
Variation of OLB mortality (proportion of dead OLB insects) among oak species in the non-choice experiment. Small symbols represent raw data, large symbols and error bars represent raw means ± SE. The gray shading area indicates that *Q. ilex* was not included in the GLMM. Different letters (a and b) indicate significant differences between oak species, whereas the asterisks indicate significant differences between herbivory treatments (damaged *vs* intact leaves). Significant differences were obtained with the post-hoc test, and were the same as in the LMM with only the three other oak species

## References

Abdala-Roberts L, Pérez Niño B, Moreira X, et al (2019) Effects of early-season insect herbivory on subsequent pathogen infection and ant abundance on wild cotton (*Gossypium hirsutum*). J Ecol 107:1518–1529. https://doi.org/10.1111/1365-2745.13131

Ali GJ, Agrawal AA (2014) Asymmetry of plant-mediated interactions between specialist aphids and caterpillars on. Funct Ecol 28:1404–1412

Bălăcenoiu F, Japelj A, Bernardinelli I, et al (2021) *Corythucha arcuata* (Say, 1832) (Hemiptera, Tingidae) in its invasive range in Europe: perception, knowledge and willingness to act in foresters and citizens. NeoBiota 69:133–153

Bates D, Maechler M, Bolker B, Walker S (2018) lme4: Linear Mixed-Effects Models using “Eigen” and S4

Bernardinelli I (2006) Potential host plants of *Corythucha arcuata* (Het., Tingidae) in Europe: A laboratory study. J Appl Entomol 130:480–484. https://doi.org/10.1111/j.1439-0418.2006.01098.x

Bernardinelli I (2000) Distribution of the Oak lace bug *Corythucha arcuata* (Say) in northern Italy (Heteroptera Tingidae). Redia 83:157–162

Bernardinelli I, Zandigiacomo P (2000) *Corythucha arcuata* (Say): a new pest for European oaks. Methodology of forest insect and disease survey in Central Europe. Proceedings of the IUFRO working party 7.03.10 workshop. September 24–28. In: Knizek M, Forster B, Grodzki W, et al. (eds) Lux Libris. Buşteni, Romania. Braşov, Romania, pp 121–122

Bernaschini ML, Valladares G, Salvo A (2020) Edge effects on insect–plant food webs: assessing the influence of geographical orientation and microclimatic conditions. Ecol Entomol 45:806–820. https://doi.org/10.1111/een.12854

Biere A, Goverse A (2016) Plant-mediated systemic interactions between pathogens, parasitic nematodes, and herbivores above- and belowground. Annu Rev Phytopathol 54:499–527. https://doi.org/10.1146/annurev-phyto-080615-100245

Bingham RA, Agrawal AA (2010) Specificity and trade-offs in the induced plant defence of common milkweed *Asclepias syriaca*. J Ecol 98:1014–1022. https://doi.org/10.1111/j.1365-2745.2010.01681.x

Carmona D, Fornoni J (2013) Herbivores can select for mixed defensive strategies in plants. New Phytol 197:576–585. https://doi.org/10.1111/nph.12023

Castagneyrol B, Halder I Van, Kadiri Y, Jactel H (2021) Host-mediated, cross-generational intraspecific competition in a herbivore species. Peer Community Ecol 1–15

Csóka G, Hirka A, Mutun S, et al (2020) Spread and potential host range of the invasive oak lace bug *[Corythucha arcuata* (Say, 1832) – Heteroptera: Tingidae] in Eurasia. Agric For Entomol 22:61–74. https://doi.org/10.1111/afe.12362

Damestoy T, Brachi B, Moreira X, et al (2019) Oak genotype and phenolic compounds differently affect the performance of two insect herbivores with contrasting diet breadth. Tree Physiol 39:615–627. https://doi.org/10.1093/treephys/tpy149

Damestoy T, Moreira X, Jactel H, et al (2021) Growth and mortality of the oak processionary moth, *thaumetopoea processionea*, on two oak species: Direct and trait-mediated effects of host and neighbour species. Entomol Gen 41:13–25. https://doi.org/10.1127/entomologia/2020/1005

De Rigo D, Caudullo G (2016) Quercus Ilex in Europe: Distribution, Habitat, Usage and Threats. Eur Atlas For Tree Species 152–153

Dobreva M, Simov N, Georgiev G, et al (2013) First record of *corythucha arcuata* (Say) (Heteroptera: Tingidae) on the Balkan Peninsula. Acta Zool Bulg 65:409–412

Erb M, Meldau S, Howe GA (2012) Role of phytohormones in insect-specific plant reactions. Trends Plant Sci 17:250–259. https://doi.org/10.1016/j.tplants.2012.01.003

Fernández de Bobadilla M, Bourne ME, Bloem J, et al (2021) Insect species richness affects plant responses to multi-herbivore attack. New Phytol 231:2333–2345. https://doi.org/10.1111/nph.17228

Gandhi KJK, Herms DA (2010) Direct and indirect effects of alien insect herbivores on ecological processes and interactions in forests of eastern North America. Biol Invasions 12:389–405. https://doi.org/10.1007/s10530-009-9627-9

Ghosh E, Sasidharan A, Ode PJ, Venkatesan R (2022) Oviposition preference and performance of a specialist herbivore is modulated by natural enemies, larval odors, and immune status. J Chem Ecol 48:670–682. https://doi.org/10.1007/s10886-022-01363-5

Glazebrook J (2005) Contrasting mechanisms of defense against biotrophic and necrotrophic pathogens. Annu Rev Phytopathol 43:205–227. https://doi.org/10.1146/annurev.phyto.43.040204.135923

Gómez S, Ferrieri RA, Schueller M, Orians CM (2010) Methyl jasmonate elicits rapid changes in carbon and nitrogen dynamics in tomato. New Phytol 188:835–844. https://doi.org/10.1111/j.1469-8137.2010.03414.x

Gómez S, Orians CM, Preisser EL (2012) Exotic herbivores on a shared native host: Tissue quality after individual, simultaneous, and sequential attack. Oecologia 169:1015–1024. https://doi.org/10.1007/s00442-012-2267-2

Herms DA, McCullough DG (2014) Emerald ash borer invasion of north america: History, biology, ecology, impacts, and management. Annu Rev Entomol 59:13–30. https://doi.org/10.1146/annurev-ento-011613-162051

Hernandez-Cumplido J, Glauser G, Benrey B (2016) Cascading effects of early-season herbivory on late-season herbivores and their parasitoids. Ecology 97:1283–1297. https://doi.org/10.1890/15-1293.1

Jeffries MJ, Lawton J. (1984) Enemy free space and the structure of ecological communities. Biol J Linn Soc 23.4:269–286

Kaplan I, Denno RF (2007) Interspecific interactions in phytophagous insects revisited: A quantitative assessment of competition theory. Ecol Lett 10:977–994. https://doi.org/10.1111/j.1461-0248.2007.01093.x

Kinahan IG, Baranowski AK, Whitney ER, et al (2020) Facilitation between invasive herbivores: hemlock woolly adelgid increases gypsy moth preference for and performance on eastern hemlock. Ecol Entomol 45:416–422. https://doi.org/10.1111/een.12829

Kovač M, Gorczak M, Wrzosek M, et al (2020) Identification of entomopathogenic fungi as naturally occurring enemies of the invasive oak lace bug, *Corythucha arcuata* (Say) (Hemiptera: Tingidae). Insects 11:679

Kroes A, Van Loon JJA, Dicke M (2015) Density-dependent interference of aphids with caterpillar-induced defenses in arabidopsis: Involvement of phytohormones and transcription factors. Plant Cell Physiol 56:98–106. https://doi.org/10.1093/pcp/pcu150

Leitner M, Boland W, Mithöfer A (2005) Direct and indirect defences induced by piercing-sucking and chewing herbivores in *Medicago truncatula*. New Phytol 167:597–606. https://doi.org/i:10.1111/j.1469-8137.2005.01426.x

Marković C, Dobrosavljević J, Milanović S (2021) Factors influencing the oak lace bug (Hemiptera: Tingidae) behavior on oaks: Feeding preference does not mean better performance? J Econ Entomol 114:2051–2059. https://doi.org/10.1093/jee/toab148

Mattson WJ (1980) Herbivory in relation to plant nitrogen content. Annu Rev Ecol Syst 11:119–161

McArt SH, Thaler JS (2013) Plant genotypic diversity reduces the rate of consumer resource utilization. Proc R Soc B Biol Sci 280:. https://doi.org/10.1098/rspb.2013.0639

Milanović S, Lazarević J, Popović Z, et al (2014) Preference and performance of the larvae of *Lymantria dispar* (Lepidoptera: Lymantriidae) on three species of european oaks. Eur J Entomol 111:371–378. https://doi.org/10.14411/eje.2014.039

Moreira X, Abdala-Roberts L, Castagneyrol B (2018) Interactions between plant defence signalling pathways: Evidence from bioassays with insect herbivores and plant pathogens. J Ecol 106:2353–2364. https://doi.org/10.1111/1365-2745.12987

Moreira X, Abdala-Roberts L, Hernández-Cumplido J, et al (2015) Specificity of induced defenses, growth, and reproduction in lima bean (*Phaseolus lunatus*) in response to multispecies herbivory. Am J Bot 102:1300–1308. https://doi.org/10.3732/ajb.1500255

Moreira X, Lundborg L, Zas R, et al (2013) Inducibility of chemical defences by two chewing insect herbivores in pine trees is specific to targeted plant tissue, particular herbivore and defensive trait. Phytochemistry 94:113–122. https://doi.org/10.1016/j.phytochem.2013.05.008

Moreira X, Zas R, Sampedro L (2012) Genetic variation and phenotypic plasticity of nutrient re-allocation and increased fine root production as putative tolerance mechanisms inducible by methyl jasmonate in pine trees. J Ecol 100:810–820

Nakagawa S, Schielzeth H (2013) A general and simple method for obtaining R2 from generalized linear mixed-effects models. Methods Ecol Evol 4:133–142. https://doi.org/10.1111/j.2041-210x.2012.00261.x

Newingham BA, Callaway RM, BassiriRad H (2007) Allocating nitrogen away from a herbivore: A novel compensatory response to root herbivory. Oecologia 153:913–920. https://doi.org/10.1007/s00442-007-0791-2

Ohgushi T (2008) Herbivore-induced indirect interaction webs on terrestrial plants: the importance of non-trophic, indirect, and facilitative interactions. Entomol Exp Appl 128:217–229

Paulin M, Hirka A, Csepelényi M, et al (2021) Overwintering mortality of the oak lace bug (*Corythucha arcuata*) in Hungary – a field survey. Cent Eur For J 67:108–112. https://doi.org/10.2478/forj-2020-0028

Paulin M, Hirka A, Eötvös CB, et al (2020) Known and predicted impacts of the invasive oak lace bug (*Corythucha arcuata*) in European oak ecosystems-A review. Folia Oecologica 47:131–139. https://doi.org/10.2478/foecol-2020-0015

Paulin M, Hirka A, Miko A, et al (2019) Tölgycsipkéspoloska – Helyzetjelentés 2019 őszén [The oak-lace bug – status report in autumn 2019]. In Alföldi Erdőkért Egyesület Kutatói Nap. Tudományos eredmények a gyakorlatban. Lakitelek 2019.

Poelman EH, Broekgaarden C, Van Loon JJA, Dicke M (2008) Early season herbivore differentially affects plant defence responses to subsequently colonizing herbivores and their abundance in the field. Mol Ecol 17:3352–3365. https://doi.org/10.1111/j.1365-294X.2008.03838.x

Poelman EH, Dicke M (2014) Plant-mediated Interactions Among Insects within a Community Ecological Perspective. Annu Plant Rev Insect-Plant Interact 47:309–337. https://doi.org/10.1002/9781118829783.ch9

R Core Team (2020) R: A Language and environment for statistical computing

Rigsby CM, Body MJA, May A, et al (2021) Impact of chronic stylet-feeder infestation on folivore-induced signaling and defenses in a conifer. Tree Physiol 41:416–427. https://doi.org/10.1093/treephys/tpaa136

Rytteri S, Kuussaari M, Saastamoinen M (2021) Microclimatic variability buffers butterfly populations against increased mortality caused by phenological asynchrony between larvae and their host plants. Oikos 130:753–765. https://doi.org/10.1111/oik.07653

Schaeffer RN, Wang Z, Thornber CS, et al (2018) Two invasive herbivores on a shared host: patterns and consequences of phytohormone induction. Oecologia 186:973–982. https://doi.org/10.1007/s00442-018-4063-0

Schweiger R, Heise AM, Persicke M, Müller C (2014) Interactions between the jasmonic and salicylic acid pathway modulate the plant metabolome and affect herbivores of different feeding types. Plant, Cell Environ 37:1574–1585. https://doi.org/10.1111/pce.12257

Singer MS, Rodrigues D, Stireman III J., Carrière Y (2004) Roles of food quality and enemy-free space in host use by a generalist insect herbivore. Ecology 85:2747–2753

Sönmez E, Demirbağ Z, Demır İ (2016) Pathogenicity of selected entomopathogenic fungal isolates against the oak lace bug, *Corythucha arcuata* Say. (Hemiptera: Tingidae), under controlled conditions. Turkish J Agric For 40:715–722. https://doi.org/10.3906/tar-1412-10

Stewart JE, Maclean IMD, Edney AJ, et al (2021) Microclimate and resource quality determine resource use in a range-expanding herbivore. Biol Lett 17:. https://doi.org/10.1098/rsbl.2021.0175

Thaler JS, Humphrey PT, Whiteman NK (2012) Evolution of jasmonate and salicylate signal crosstalk. Trends Plant Sci 17:260–270. https://doi.org/10.1016/j.tplants.2012.02.010

Van Dijk LJA, Ehrlén J, Tack AJM (2020) The timing and asymmetry of plant-pathogen-insect interactions: Plant-pathogen-insect interactions. Proc R Soc B Biol Sci 287:. https://doi.org/10.1098/rspb.2020.1303

Wetzel WC, Kharouba HM, Robinson M, et al (2016) Variability in plant nutrients reduces insect herbivore performance. Nature 539:425–427. https://doi.org/10.1038/nature20140

